# Theta oscillations optimize a speed-precision trade-off in phase coding neurons

**DOI:** 10.1101/2022.12.08.519523

**Authors:** Adrián F. Amil, Albert Albesa-González, Paul F.M.J. Verschure

## Abstract

Low-frequency oscillations shape how neurons sample their synaptic inputs, regulating information exchange across networks. In the hippocampus, theta-band oscillations (3–8 Hz) reorganize cortical input signals temporally, resulting in a phase code. However, the reason hippocampal oscillations are limited to low frequencies like the theta band remains unclear. Here, we derive a theoretical framework for neuronal phase coding to show that realistic noise levels create a trade-off between sampling speed (controlled by oscillation frequency) and encoding precision in hippocampal neurons. This speed-precision trade-off produces a maximum in information rate within the theta band of *~*1–2 bits/s. Additionally, we demonstrate that our framework explains other key hippocampal properties, such as the preservation of theta along the dorsoventral axis despite various physiological gradients, and the modulation of theta frequency and amplitude by the animal’s running speed. Extending our analysis to extra-hippocampal areas, we propose that theta oscillations may also support efficient encoding of stimuli in visual cortex and olfactory bulb. More broadly, we lay the groundwork for rigorously studying how system constraints determine optimal sampling frequency regimes for phase coding neurons in biological and artificial brains.

**Author Summary:** The rodent hippocampus exhibits prominent oscillations in the theta band (3–8 Hz) during exploration, enabling individual neurons to rhythmically sample and represent sensory signals from the cortex. However, the reason behind the specific frequency of this hippocampal rhythm has remained unclear. In this study, we developed a biologically-based theoretical framework to demonstrate that neurons using oscillations to efficiently sample noisy signals encounter a trade-off between their sampling speed (i.e., oscillation frequency) and their coding precision (i.e., reliability of encoding). Notably, our findings reveal that this trade-off is optimized precisely within the theta band, while also providing insights into other fundamental features. In conclusion, we offer an explanation grounded in efficient coding for why hippocampal oscillations are confined to the theta band and establish a foundation for exploring how the properties of individual neurons determine optimal sampling frequencies in specific neural circuits.

## 1 Introduction

Early physiological experiments with rodents revealed that certain neurons exhibit increased firing rates when the animal occupies specific regions within its environment, referred to as place fields (O’Keefe, 1976). Further research revealed that these place cells also exhibit a temporal pattern known as phase precession: as the animal traverses a place field, they progressively fire at earlier phases of the local field potential (LFP) oscillations (O’Keefe and Recce, 1993). This discovery spurred extensive research into the relationship between behavior and the timing of neuronal firing relative to theta-band (3–8 Hz) oscillations. It was found that incorporating firing phase information alongside firing rates significantly enhanced the accuracy of reconstructing the animal’s position (Jensen and Lisman, 2000), establishing the phase code as a viable alternative to the traditional firing rate code in neural circuits (Lisman, 2005).

The phase precession phenomenon also led to the development of theories and models that would explain the temporal progression of spikes across the theta phase. One influential model is the rate-to-phase transform, where firing rate input from the entorhinal cortex make hippocampal neurons to fire earlier or later depending on firing rate levels, transforming rate-coded inputs into phase-coded outputs (McLelland and Paulsen, 2009; Mehta, A. Lee, and Wilson, 2002). Empirical evidence supporting this model has been observed also outside the hippocampus (Margrie and Schaefer, 2003), suggesting a broader application to sensory-related areas (Lisman, 2005). Indeed, the rate-to-phase transform can be linked to synaptic sampling (Buesing et al., 2011), where low-frequency field oscillations modulate neuronal excitability –via ephaptic effects (Anastassiou et al., 2011) or indirectly via feedback inhibition (Fries, 2015; Mizuseki et al., 2009). By leveraging a rhythmically-organized first-spike latency code (VanRullen, Guyonneau, and Thorpe, 2005), phase coding can sample rate-based synaptic inputs each cycle. This rhythmic neural sampling from sensory-related areas would then manifest as perceptual sampling effects at the behavioral level (Caplette, Jerbi, and Gosselin, 2022; Korcsak-Gorzo et al., 2022; VanRullen, 2016). Behavioral and electrophysiological experiments support this hypothesis, showing that stimulus detection is influenced by the timing of stimulus presentation relative to the oscillatory cycle in visual (Kasten and Herrmann, 2020) and olfactory (Kepecs, Uchida, and Mainen, 2006; Smear et al., 2011) domains. Furthermore, the idea of rhythmic neural sampling aligns with the short membrane time constants of pyramidal cells (10-30 ms) compared to the periods of low-frequency field oscillations (100 ms to 1 s), allowing for cycle-to-cycle resets. Therefore, rhythmic field activity along the sensory hierarchy could provide a scaffold for representing sensory signals through repeated synaptic input sampling, suggesting that rhythmic sampling might also facilitate probabilistic inference of the underlying sensory streams (Buesing et al., 2011; Korcsak-Gorzo et al., 2022).

Following the efficient coding hypothesis (Barlow et al., 1961), oscillations in brain circuits may have been evolutionarily optimized for information encoding and transmission. The crucial role of oscillations in neuronal communication (Fries, 2015) and the conservation of basic oscillatory rhythms across species (Buzsáki, Logothetis, and Singer, 2013) support this hypothesis. However, a fundamental question remains: why has this low-frequency range emerged over others that could offer increased bandwidth and sampling resolution? In this paper, we address why oscillations supporting phase coding seem to be confined to low frequencies like the theta band. Through theoretical analysis and simulations, we show that theta-band frequencies strike a balance between sampling speed (i.e., oscillation frequency) and encoding precision (i.e., information per oscillation cycle), resulting in an optimal information rate. We further extend our analysis to account for the persistence of theta oscillations along the hippocampal dorsoventral axis, their modulation by the animal’s running speed, and also their presence in extra-hippocampal brain areas like the primary visual cortex and the olfactory bulb. Finally, we discuss how our approach can contribute towards understanding oscillations through the lenses of optimality and efficient coding.

## 2 Theoretical framework

We first introduce our theoretical framework, where we derive and validate an analytical approximation of the information rate conveyed by phase coding neurons.

### 2.1 Neuron model

We model pyramidal cells as stochastic leaky integrate-and-fire (LIF) neurons receiving oscillatory and tonic inputs (Burkitt, 2006) (Figure 1). Their membrane potential dynamics are defined by:

**Figure 1:**
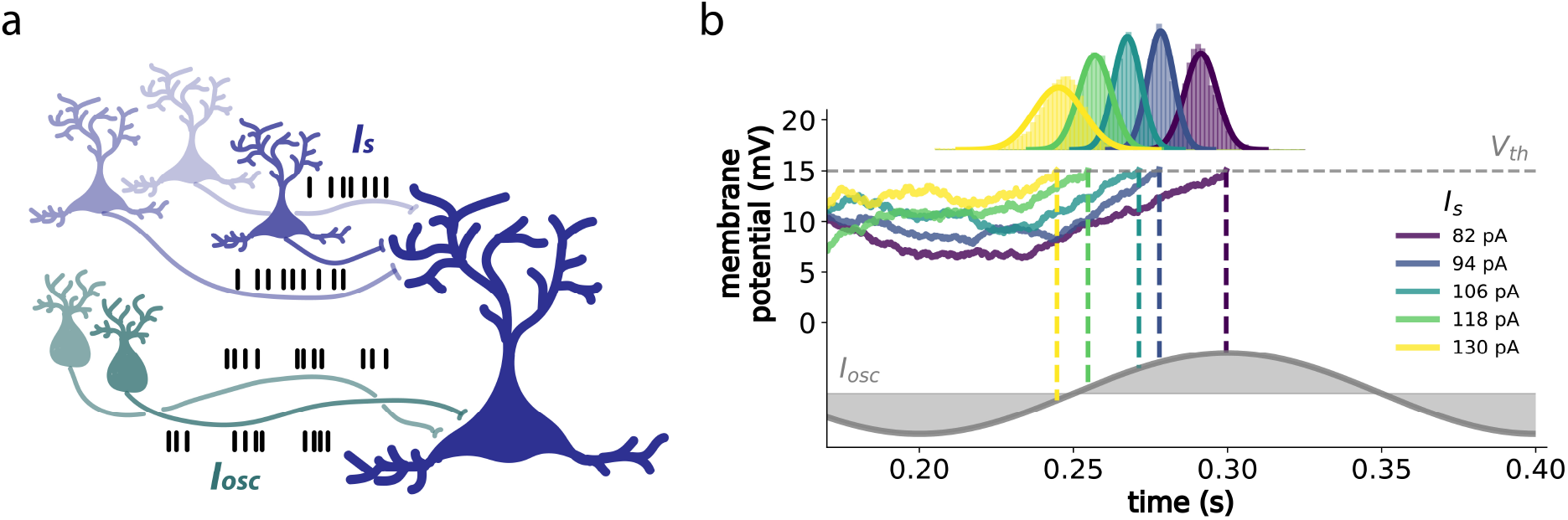
Phase coding in LIF neurons. (a) Model neuron receiving rhythmic volleys of inhibitory spikes to the soma, and continuous volleys of excitatory spikes to the dendrites, representing the oscillatory and tonic inputs in Equation 1, respectively. (b) Examples of voltage trajectories for such a neuron with hippocampal parameters (McLelland and Paulsen, 2009) and realistic noise amplitudes (Lansky, Sanda, and He, 2006) (see Table 1 in Appendix 5.4 for details on parameters), for a range of tonic inputs *I*_*s*_ (color code). Vertical dashed lines denote the times at which the neuron reaches threshold (denoted by the horizontal dashed line), with respect to the oscillation (i.e., phase). Histograms and the corresponding Gaussian fits denote the probability distributions of spike times for each *I*_*s*_.

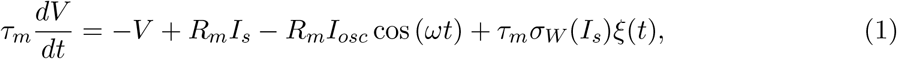

with *V* (*t*′ + *dt*) = 0 if *V* (*t*′) ≥ *V*_*th*_. Here, *V* is the membrane potential, *R*_*m*_ the membrane resistance, *τ*_*m*_ the membrane time constant, *I*_*s*_ the tonic input current that conveys the stimulus information to be encoded, *I*_*osc*_ the oscillation amplitude, *ω* = 2*πf* the angular frequency, *V*_*th*_ the spike threshold, *σ*_*W*_ (*I*_*s*_) the signal-deependent noise amplitude, and *ξ*(*t*) ≡ *dW* (*t*)*/dt* the Wiener process derivative with 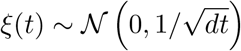.

#### Noise

We assume input currents in our neuron model have a synaptic origin (Figure 1a). A high-rate Poisson input with synaptic time constants *τ*_*s*_ ≪ *τ*_*m*_ leads to Gaussian white noise, *ξ*(*t*), in Equation 1. Although biological neurons display colored noise (Brunel et al., 2001; Demir, 2002; Schwalger, Droste, and Lindner, 2015), Gaussian white noise is still a reasonable approximation (see Burkitt, 2006; Lansky, Sanda, and He, 2006) and is analytically tractable. Under these assumptions, the standard deviation of the free membrane potential (i.e., the membrane potential without considering the threshold *V*_*th*_), denoted as *σ*_*V*_, is directly proportional to the noise strength *σ*_*W*_. In turn, it can be demonstrated that *σ*_*V*_ is proportional to 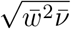, where 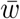 represents the mean synaptic efficacy, and 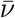 denotes the mean presynaptic firing rate (Korcsak-Gorzo et al., 2022). Defining the effective tonic input as 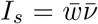, it then follows that 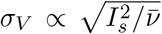. Importantly, due to the membrane’s low-pass filtering properties (*τ*_*m*_), *I*_*s*_ must increase with oscillation frequency *f* to maintain phase-locking (Figure S1c, d). We assume these adjustments result from changes in 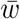, keeping 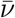 constant across frequencies. Hence, we incorporate signal-dependent noise from Poisson processes (where variance equals the mean) by scaling the noise amplitude *σ*_*W*_ by *KI*_*s*_, where *K* is a constant representing the inverse of *I*_*s,f*=1_, the weakest *I*_*s*_ possible that makes the neuron phase-lock to a 1 Hz oscillation—acting as a baseline. Therefore, *σ*_*W*_ can be defined as:

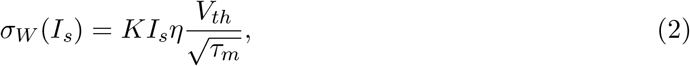

with *KI*_*s*_ as the signal-dependent factor akin to the linear modulation of noise amplitude in motor control (Harris and Wolpert, 1998; Jones, Hamilton, and Wolpert, 2002), *η* a dimensionless parameter controlling noise strength, and 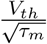 scaling *σ*_*W*_ to the noise units 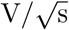.

### 2.2 Approximation of spike phase distributions

Given the neuron model described by Equations 1 and 2, we can proceed to derive an analytical approximation for the mean and variance of the spike phase distributions (e.g., Figure 1b).

#### Mean phase

As demonstrated in previous work (McLelland and Paulsen, 2009), we can derive a closed-form expression for the expected phase *µ*_*ϕ*_ at which the neuron described by Equation 1 will phase-lock. By integrating the deterministic part of Equation 1, we can find the phase at which the expected trajectory of the membrane potential *µ*_*V*_ (*t*) intersects with the spike threshold *V*_*th*_, exactly after one oscillation period *T* (see the Appendix 5.1 for the full derivation), obtaining:

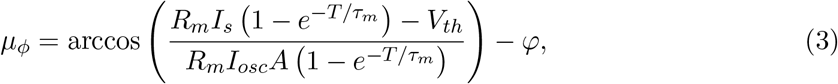

where 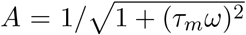, representing the filtering of the oscillatory input by the membrane, and *φ* = − arctan (*ωτ*_*m*_), representing a phase shift. Equation 3 captures how the integrated tonic and oscillatory inputs make a neuron reach threshold *V*_*th*_ with a period *T* between spikes (see Figure S1a–c for details).

#### Phase variance

To estimate the variance of spike phase 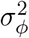 due to noise, we consider the variance in spike timing 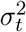 induced by fluctuations in the membrane potential. Similarly to previous work on phase jitter in oscillators (Demir and Sangiovanni-Vincentelli, 2012; Kilinc and Demir, 2018), we employ a first-order Taylor approximation of the membrane dynamics around the spike threshold *V*_*th*_, such that 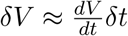, and consider the integrated noise 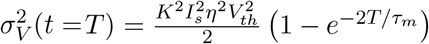. Using the propagation of uncertainty, we express 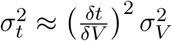. Then, since time and phase are related by *ϕ* = *ωt*, we have that 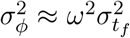. Hence, we arrive at the following approximation (see Figure 2a, and Appendix 5.2 for more details):

**Figure 2:**
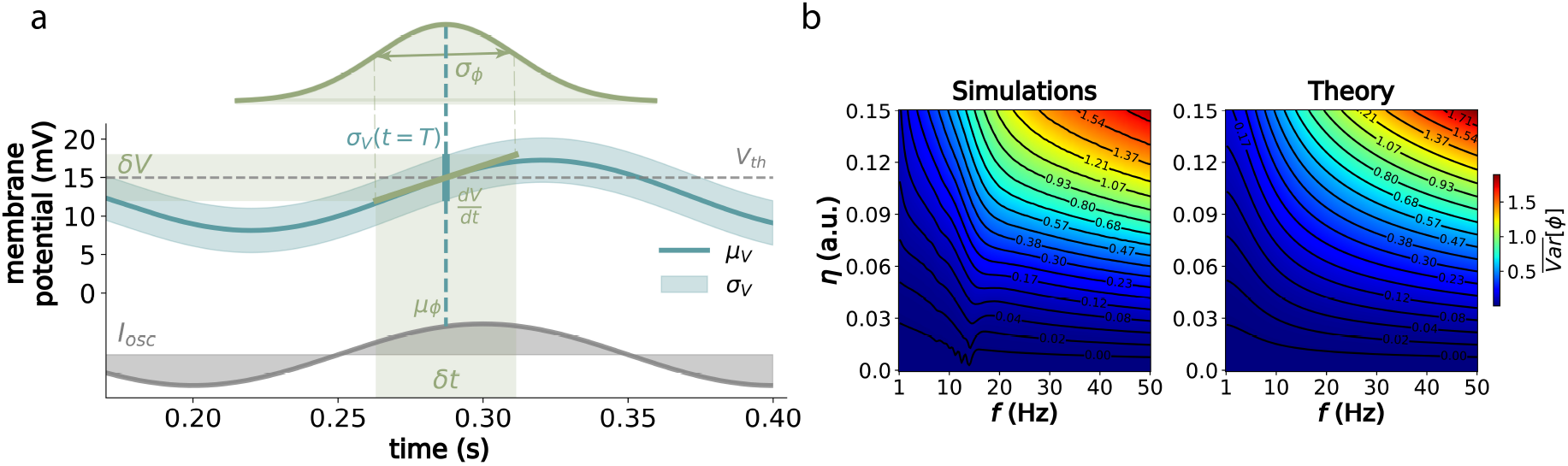
Analytical approximation of phase distributions. (a) Approximation of varian1ce in phase of firing depends on the interplay between the linearized membrane dynamics at spike threshold 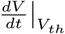 (with *µ*_*V*_ being the free membrane potential) and the accumulated noise in voltage *σ*_*V*_ (*t*) at the expected firing time after one period *T*, so that *δV* ≡ *σ*_*V*_ (*t* = *T*) and *δt* ≡ *σ*_*t*_. The probability distribution on the upper part denotes how *δt* translates to *σ*_*ϕ*_ with respect to the oscillation (lower part). For this example, we used the same parameters as in Figure 1, defined in Table 1 in Appendix 5.4, for an *I*_*s*_ of 90 pA. (b) Mean variance (in rad^2^) across *I*_*s*_ levels for physiologically-relevant parameters of frequency and noise, for both simulations and the analytical predictions from Equation 4.

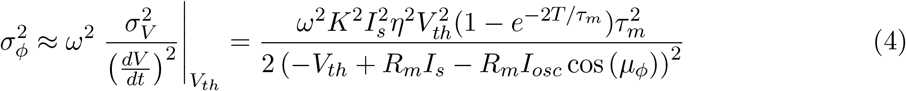

Equation 4 describes how variance in spike phase results from the interaction between the accumulated random fluctuations in membrane potential and the membrane dynamics near the spike threshold. This linear approximation is valid for small deviations from the threshold (low noise) and a nearly linear response around it (suprathreshold regime). Notably, it agrees well with simulations (Figure 2b), predicting 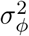 accurately under physiologically-relevant parameters and noise levels (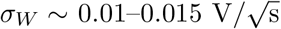 in Lansky, Sanda, and He (2006) and McLelland and Paulsen (2009), corresponding to *η ~* 0.1–0.15 in Figure 2b). The predictions align with simulations until noise levels become unrealistically high (*η* > 0.2) and primarily at oscillation frequencies above 30 Hz (Figure S2). These deviations are largely due to variance being unbounded in theory (Equation 4) but bounded to *~ π*^2^*/*3 rad in simulations (i.e., with phase becoming approximately uniformly distributed in [0, 2*π*]). Moreover, in the regime of very high variance, these differences are negligible considering the limited range of [0, 2*π*], having a minimal impact on information estimates. Thus, we conclude that phase can be generally well characterized by a normal distribution with mean *µ*_*ϕ*_ (Equation 3) and variance 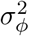 (Equation 4).

### 2.3 Approximation of information rate

Using the mean and variance of the phase distributions, we can approximate the neuron’s information about the stimulus *I*_*s*_ per cycle. Assuming *I*_*s*_ is uniformly distributed across the phase-locking range (with *M* equally-spaced *I*_*s*_), the phase distribution forms a Gaussian mixture with *M* means ***µ*** and variances ***σ***^2^. The mutual information between *I*_*s*_ and *ϕ* can be approximated by considering an aggregate measure of the overall spread of the mixture (see Appendix 5.3 for details), obtaining:

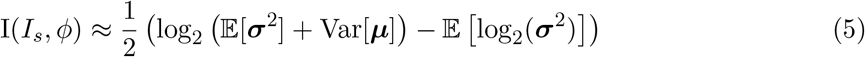

This equation provides a good approximation of the information a spike phase contains about *I*_*s*_ at every cycle (Figure 3).

**Figure 3:**
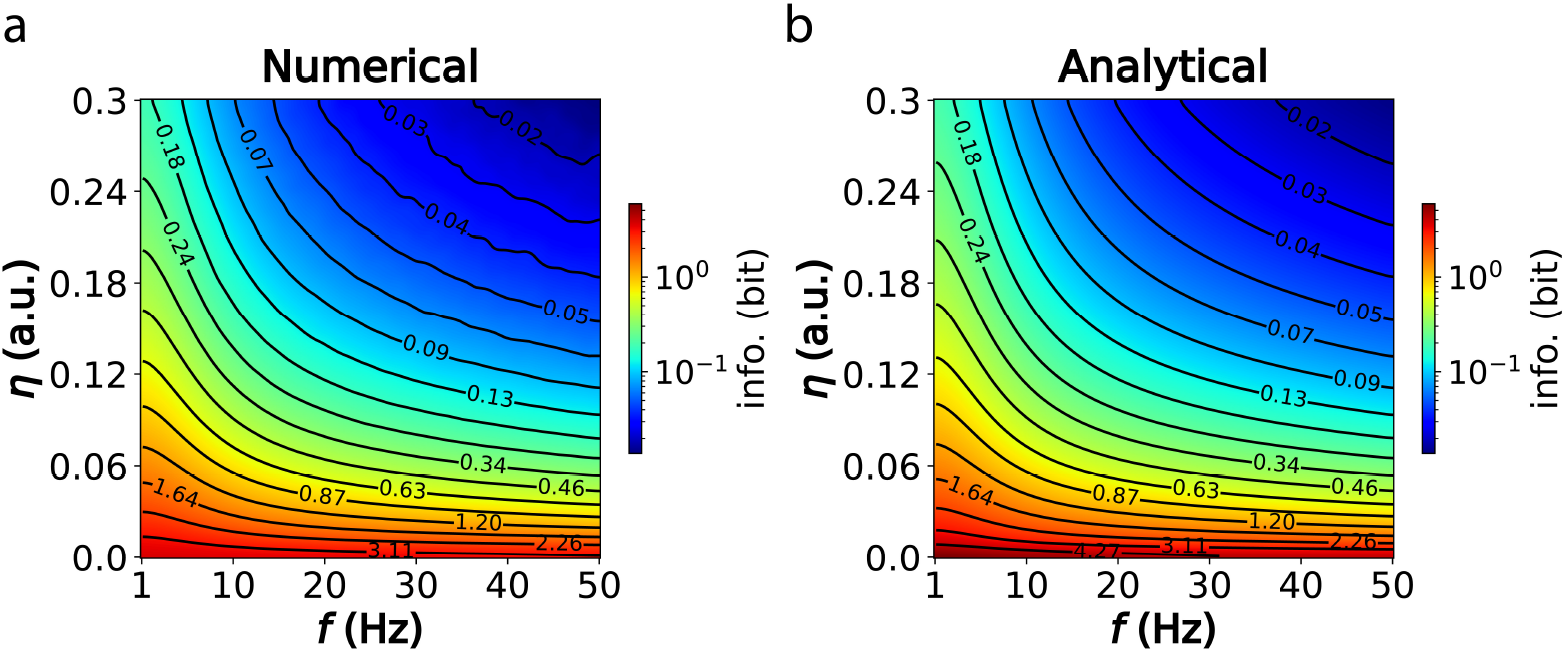
Approximation of the mutual information. (a) Mutual information via numerical integration of the Gaussian mixture. For each point in the frequency-noise parameter space, we estimated the means and variances of the phase distributions. We then drew 10000 samples per Gaussian component to construct a histogram of 100 bins, from which entropy was computed using the Shannon formula 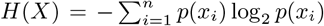. (b) Analytical approximation using Equation 5. The color map is re-scaled to log_10_ for improved visualization.

The efficient coding hypothesis (Barlow et al., 1961) suggests that neurons maximize information transfer, minimizing redundancy. Viewing oscillations as rhythmic input sampling (VanRullen, 2016), we then cast the problem of encoding a signal *s* in terms of maximizing information rate (bit/s). Simply multiplying information per cycle and oscillation frequency *f* gives r_*upper*_ ≈ I*f*, an upper bound that assumes cycle independence. However, a more realistic approach needs to account for cross-cycle correlations due to the relationship between the input signal autocorrelation 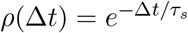 (with characteristic time constant *τ*_*s*_) and the oscillation period *T*, hence modifying the information rate to (more details in Appendix 5.3):

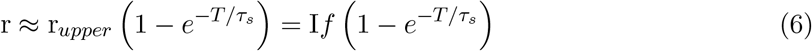

This equation reflects the impact of temporal correlations, penalizing oversampling when *T* < *τ*_*s*_ by introducing an effective frequency factor (Figure S3). Since we are mainly interested in finding the optimal frequency for given neuron parameters and noise level, we define the normalized information rate (r_*norm*_) as the ratio of r at a specific frequency (r_*f*_) to the maximum r across all frequencies (*F*):

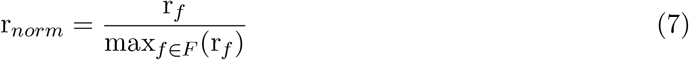

Hence, this metric allows us to study the encoding capacity of neurons across the biologically relevant spectrum of oscillation frequencies.

## 3 Results

### 3.1 The speed-precision trade-off

Maximizing encoding precision requires operating at physiologically low frequencies to mitigate noise effects and enhance signal fidelity (Figure 4a). However, this approach inherently reduces the rate of input sampling, potentially resulting in delayed reaction times. In dynamic environments, where rapid responses are essential for survival, animals cannot afford to prioritize precision at the expense of speed. Conversely, operating at excessively high frequencies may lead to redundant input sampling and an amplification of noise (Figure 4a). Therefore, neural circuits must strike a balance that allows for both precise sensory information processing and sufficiently fast reaction times. This balance reflects a fundamental speed-precision trade-off faced by animals with inherently noisy brains (see Lahiri, Sohl-Dickstein, and Ganguli, 2016 for a broader discussion).

**Figure 4:**
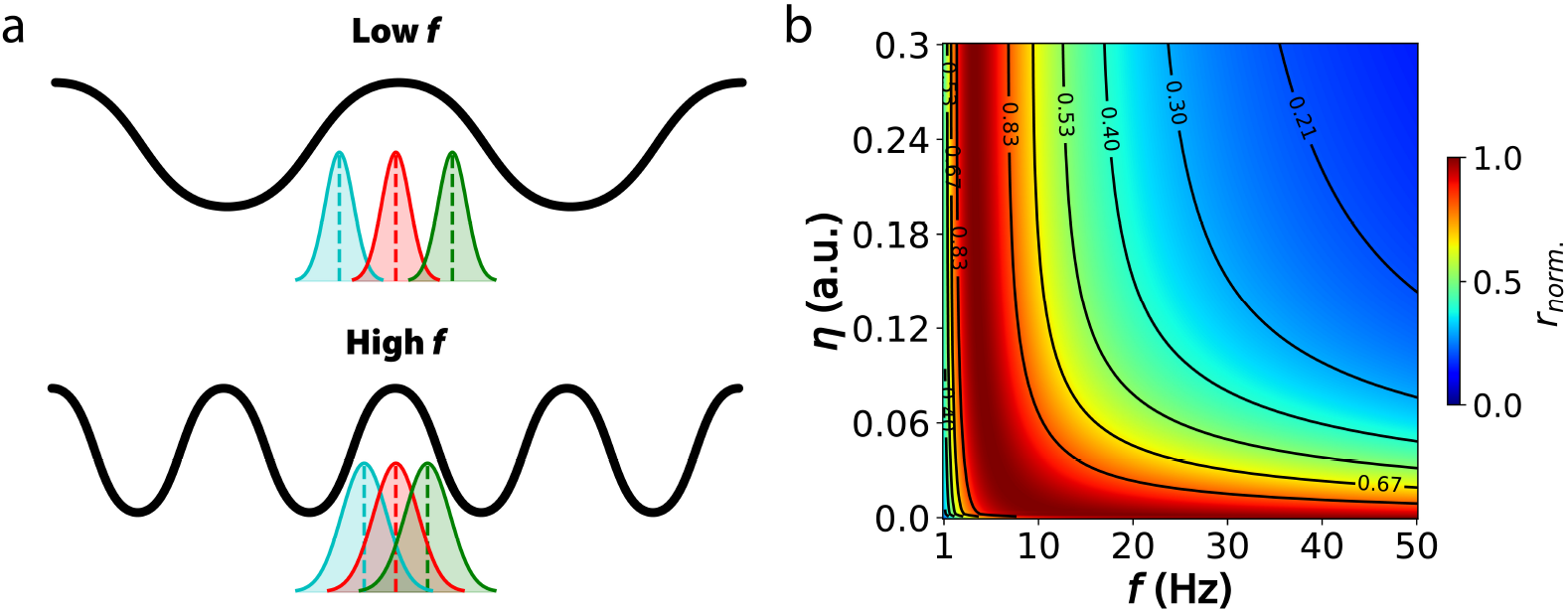
Information rate is maximized in the theta band for hippocampal neurons. (a) Implications for signal encoding at lower versus higher oscillation frequencies. (b) Information rate (bit/s, normalized across frequencies) for hippocampal neurons across a wide range of oscillation frequencies and noise strength levels. Neuron and oscillation parameters are detailed in Table 1 in Appendix 5.4. The input characteristic time constant *τ*_*s*_ was set to 100 ms. Physiologically-realistic noise strength levels correspond to the range *η ~* 0.1–0.15 (Lansky, Sanda, and He, 2006).

The concept of information rate introduced earlier—defined as the expected number of bits transmitted per second—serves as a useful metric for capturing this trade-off, as it incorporates both the precision of encoded information and the speed of processing. In our theoretical model, we investigated this sampling-precision trade-off using physiologically realistic parameters of hippocampal neurons driven at various oscillatory frequencies and noise levels. Notably, we observed a peak in the information rate within the theta frequency range (3-8 Hz) across almost the entire noise spectrum (Figure 4b), corresponding to *~*1–2 bits/s for physiologically-realistic noise levels (Figure S4). At very low noise levels, higher frequencies do not amplify noise sufficiently to significantly degrade information, thus remaining optimal. However, even a small increase in noise shifts the optimal frequency range to lower frequencies (*~*2–10 Hz). We further validated these theoretical findings with simulations (Figure S5). Additionally, we found that the lower frequency range remains optimal across a broad range of input signal time constants (Figure S6), lending further support to our results.

### 3.2 Theta optimality along the hippocampal dorsoventral axis

The hippocampus exhibits consistent variation in neuronal and circuit properties along its dorsoventral (DV) axis (Soltesz and Losonczy, 2018; Witter et al., 2000). Specifically, features such as membrane input resistance, action potential threshold, membrane time constant, and oscillation amplitude change nearly linearly along this axis (Malik et al., 2016; Patel et al., 2012). Notably, despite these gradients, theta frequency remains dominant across the DV axis (Patel et al., 2012) (Figure 5a). Given this observation and our previous results, we hypothesized that the co-variation of these physiological parameters ensures that theta optimizes the speed-accuracy trade-off across the entire DV axis. Using neuron models defined by the physiological gradients along the DV axis (Malik et al., 2016), we estimated the normalized information rate across oscillation frequencies. As predicted, theta frequency consistently peaked across the entire DV axis (see Figure 5b, and Figure S7 for further validation by simulations), supporting its optimality for phase coding throughout the hippocampus.

**Figure 5:**
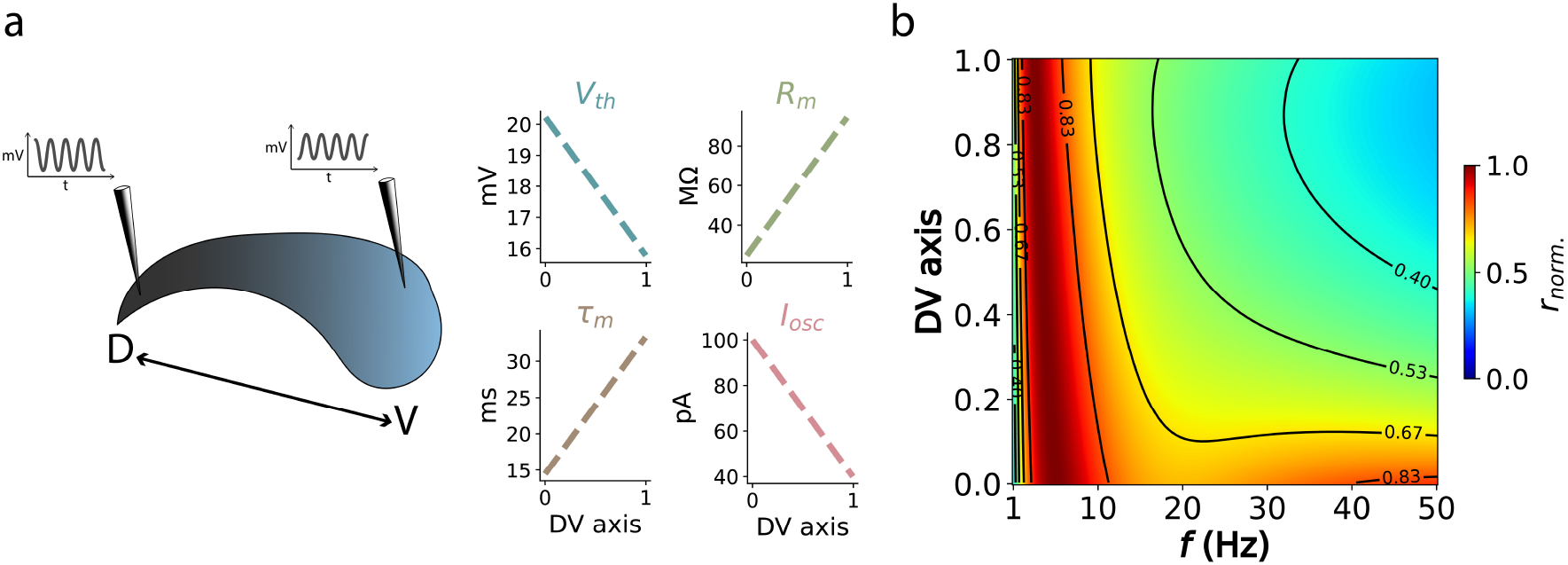
Theta remains optimal along the dorsoventral axis. (a) Physiological gradients along the DV axis of the hippocampus, extracted from Malik et al., 2016 and Patel et al., 2012. (b) Information rate for neuron models along the DV axis, across frequencies. Detailed parameters can be found in Table 2 in Appendix 5.4.

This theta optimality can be understood through the constancy of the *R*_*m*_*/τ*_*m*_ ratio, which maintains a consistent transfer function (McLelland and Paulsen, 2009). A similar relationship can be deduced by recognizing the inverse relationship between these parameters in determining the spike phase variance in Equation 4. Additionally, lower *I*_*osc*_ in ventral regions are counter-balanced by increased excitability due to lower *V*_*th*_ (Dougherty, Islam, and Johnston, 2012). In conclusion, physiological gradients along the DV axis, likely reflecting different input processing needs (M.-B. Moser and E. Moser, 1998; Witter et al., 2000), co-vary to allow optimal input sampling at theta frequency throughout the hippocampus, thereby facilitating information flow and integration across regions.

### 3.3 Optimal modulation of theta frequency and amplitude by locomotion speed

As animals move, they adjust their locomotion speed to meet task demands. Faster movement increases the rate of incoming stimuli, potentially requiring a higher frequency of perceptual sampling and corresponding brain rhythms (e.g., to normalize memory content (Pata et al., 2022)). Following this reasoning, and since theta oscillations govern the hippocampal processing of cortical inputs, it would be expected that theta frequency increases with speed (Figure 6a). Indeed, studies have shown a linear increase in theta frequency with locomotion speed in the rodent hippocampus, particularly in dorsal regions that receive sensory signals (McFarland, Teitelbaum, and Hedges, 1975; Sławińska and Kasicki, 1998; Young, Ruan, and McNaughton, 2020). This relationship has also been observed in the human posterior hippocampus (Goyal et al., 2020). Furthermore, recent analyses indicate that it is specifically speed, not acceleration, that modulates both theta amplitude and frequency (Kennedy et al., 2022). In turn, it has been suggested that hippocampal theta modulates local oscillatory circuits to maintain a consistent relationship between the rate of inputs from place field activity and spike phase (Geisler et al., 2007). Additionally, theta amplitude (and power) also exhibit a near-linear relationship with locomotion speed (Kennedy et al., 2022; Long, Bunce, and Chrobak, 2015; Young, Ruan, and McNaughton, 2020). We then hypothesized that this concurrent modulation of theta frequency and amplitude with speed might reflect an underlying objective of maximizing information rate.

**Figure 6:**
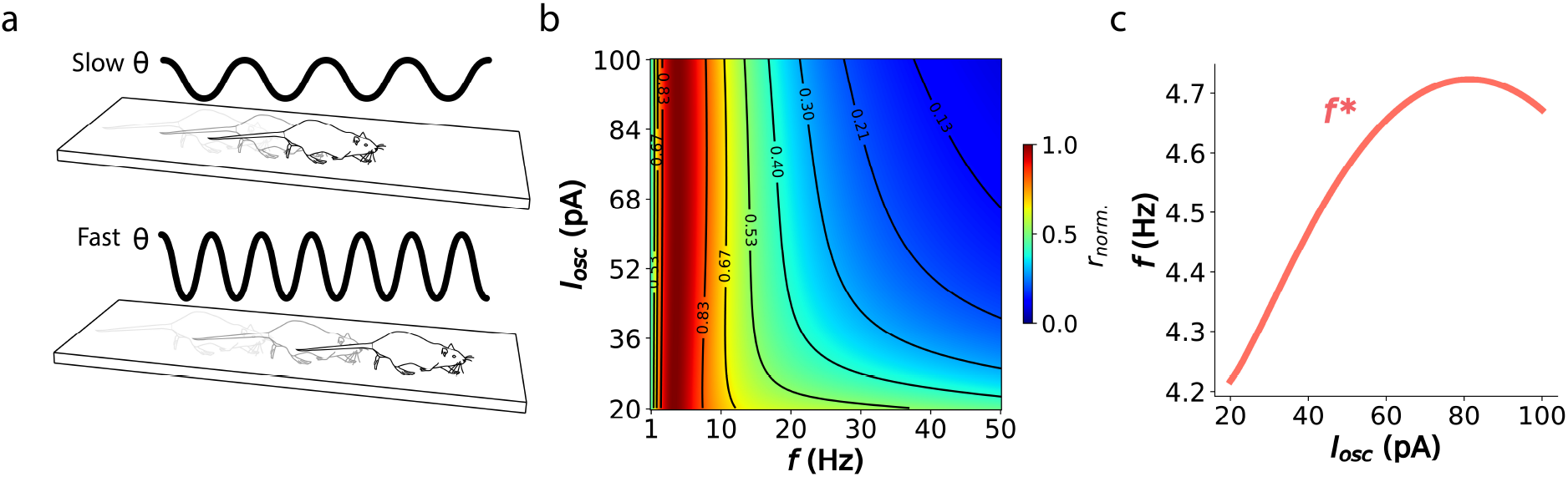
Optimal sampling requires linear modulation of theta frequency and amplitude. (a) Relationship between the rodent’s locomotion speed on a linear track and hippocampal oscillation frequency. (b) Normalized information rate across frequencies and oscillation amplitudes. Neuron parameters can be found in Table 1 in Appendix 5.4. (c) Optimal oscillation frequency defined as the information rate peak across oscillation amplitudes.

To test our hypothesis, we estimated the information rate profiles across different oscillation frequencies and amplitudes (*I*_*osc*_). Interestingly, we observed a subtle yet consistent peak shift towards higher theta frequencies as the amplitude increased (Figure 6b, c; and Figure S8 for validation by simulations). This slight shift exhibited a nearly linear relationship before reaching saturation, consistent with previous findings. Therefore, our results suggest that maximizing information rate requires the concurrent modulation of both frequency and amplitude, providing an explanation for the observed correlation between oscillatory features and the animal’s locomotion speed.

### 3.4 Generalization to extra-hippocampal areas

Our results so far suggested that the information rate maximization principle might explain the presence and adaptation of theta oscillations in the hippocampus, as efficient phase-coding neurons overcome the trade-off between speed and precision during oscillatory input sampling. However, phase coding does not seem to be unique to the hippocampus. Pyramidal cells in the primary visual cortex of rodents (Fournier et al., 2020; Huang et al., 2020; Levy et al., 2017) and monkeys (Kienitz et al., 2021; H. Lee et al., 2005; Montemurro et al., 2008; Spyropoulos, Bosman, and Fries, 2018) also exhibit theta phase locking, which has been associated with feed-forward processing of visual stimuli in superficial layers. Similarly, mitral cells in the olfactory bulb of mice are strongly modulated by theta oscillations that align with the respiratory cycle (Fukunaga et al., 2014; Losacco et al., 2020; Margrie and Schaefer, 2003).

Based on these observations, we hypothesized that low-frequency oscillations would also maximize the information rate in these sensory areas. Consistent with our hypothesis, we found a peak in the high theta-low alpha band for both models: pyramidal neurons in primary visual cortex (Figure 7a) and mitral cells in the olfactory bulb (Figure 7b).

**Figure 7:**
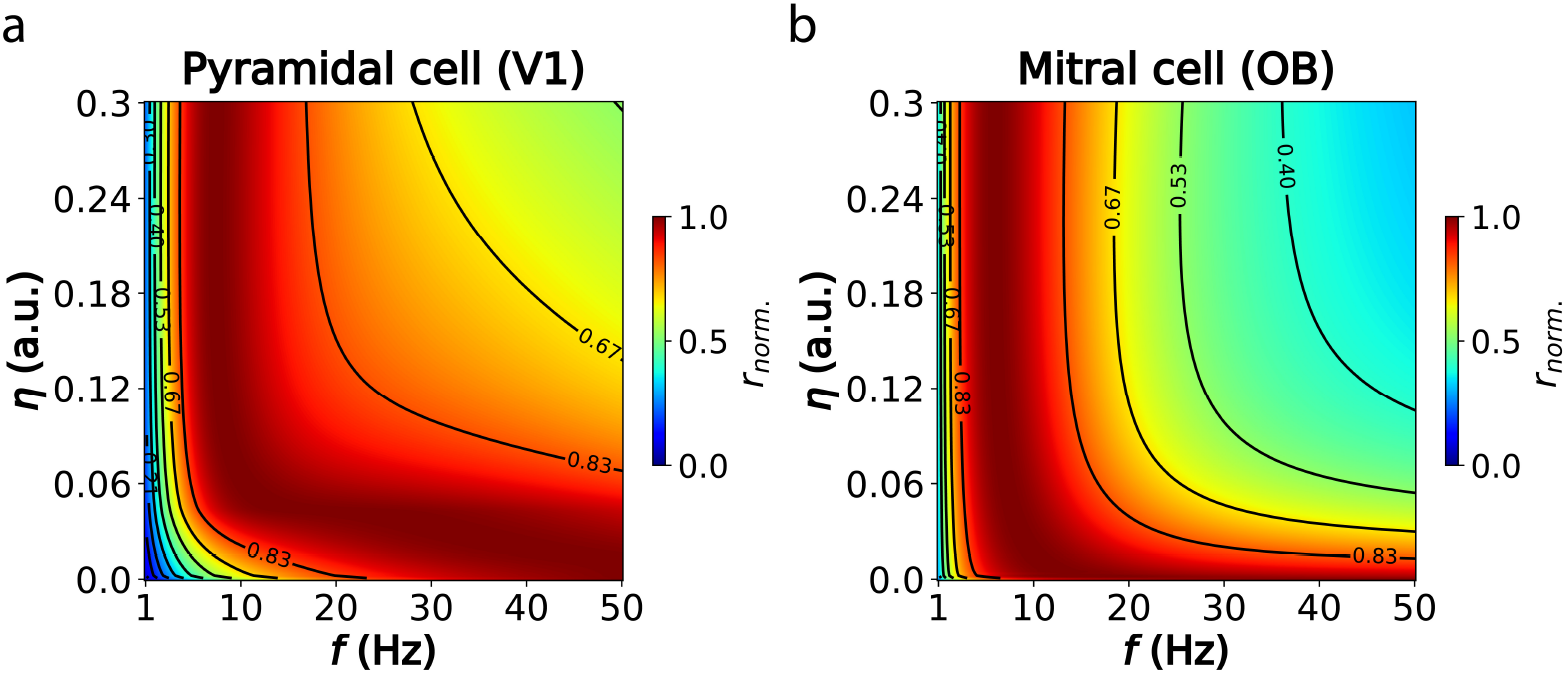
Low frequencies maximize information rate in visual cortex and olfactory bulb. (a) Normalized information rate across a broad frequency-noise parameter space for prototypical pyramidal cells in the primary visual cortex (V1) of mice (Teeter et al., 2018). A faster time constant of the input signal (*τ*_*s*_) of 50 ms was used to account for the rapid signal fluctuations in visual cortex. (b) Same as (a), but for mitral cells in the olfactory bulb of mice (Margrie and Schaefer, 2003). A slower *τ*_*s*_ of 250 ms was applied to match the cycle period of the respiratory rhythm. Detailed parameters of both neuron models can be found in Table 3 in Appendix 5.4.

## 4 Discussion

Our findings highlight the role of theta-band (3-8 Hz) oscillations in optimizing the encoding of information in phase-coding neurons, particularly within the hippocampus. By developing a theoretical framework that approximates the information rate conveyed by phase-coded signals, we have shown that theta oscillations represent an optimal solution to the trade-off between sampling speed and encoding precision. This trade-off is governed by the interplay between oscillation frequency and neuronal noise, resulting in a peak information rate within the theta frequency range.

### Relevant timescales and constraints in the hippocampus

Our results suggest that low-frequency, theta oscillations strike a balance that maximizes the information rate, enabling efficient phase coding that supports both precise sensory processing and rapid behavioral responses. Interestingly, our estimate of *~*1–2 bits/s matches the overall slowness of behavioral output rate of *~*1–10 bits/s reported in many different tasks and experimental paradigms (Zheng and Meister, 2024). In addition, the emergence of the theta range in the hippocampus as the optimal sampling frequency may impose constraints on the timescales of cortical input signals necessary for reliable encoding. Given that the bandwidth (B) of a signal with an exponentially-decaying autocorrelation function is B = 1*/*(*πτ*_*s*_), and since according to the Nyquist-Shannon theorem the sampling frequency should satisfy *f* ≥ 2B, we predict that cortical input signals (e.g., from the entorhinal cortex) should approximately follow *τ*_*s*_ ≥ 80 ms. Thus, signals with faster rates of change may risk communication losses, particularly if higher frequency ranges are employed to convey information. Therefore, we predict that rate-based input signals from the entorhinal cortex are characterized by relatively long time constants.

### Multi-modal, multi-scale integration through theta waves

Despite significant physiological gradients along the hippocampal dorsoventral (DV) axis, theta oscillations remain the dominant frequency across this axis. Our theoretical framework suggests that the co-variation of key physiological parameters optimizes the speed-precision trade-off, ensuring theta’s persistence throughout the hippocampus. We suggest that this uniform theta frequency likely facilitates the integration and transfer of information across different spatial scales and levels of resolution (Kjelstrup et al., 2008), serving as well as a consistent readout mechanism (Jensen, 2001). In turn, dorsoventral traveling waves, which follow the DV axis (Patel et al., 2012), may further enable this phase-based integration. In addition, different regions along the DV axis process inputs with varying modalities and statistics, from slow olfactory signals to fast auditory ones. The theta-based phase code might then allow local tuning of single-cell properties to diverse inputs while integrating them within a common coding format, enabling standardized and multiplexed encoding.

### Speed-controlled oscillators for optimal sampling

The linear relationship between theta frequency-amplitude and locomotion speed in rodents supports our theoretical predictions, suggesting that the hippocampus adjusts these parameters to maintain optimal information rates and efficient sensory encoding. Indeed, theta frequency fluctuations with locomotion speed (McFarland, Teitelbaum, and Hedges, 1975; Sławińska and Kasicki, 1998; Young, Ruan, and McNaughton, 2020) are driven by increased cue densities, which may require higher frequencies to match stimulus rates and normalize information per theta cycle (Pata et al., 2022), thereby helping downstream circuits maintain consistent decoding (Jensen, 2001). However, we have shown that frequency increases alone may degrade information rates due to noise; therefore, concurrent amplitude increases are necessary to sustain optimal phase coding. The medial septum, the primary driver of hippocampal theta oscillations (Buzsáki, 2002; Petsche, Stumpf, and Gogolak, 1962; Pignatelli, Beyeler, and Leinekugel, 2012), likely coordinates these adjustments, as it modulates both locomotion speed and theta frequency (Fuhrmann et al., 2015; Tsanov, 2017). Indeed, cooling experiments suggest the presence of a global oscillator in the medial septum that normalizes phase relationships by modulating theta frequency and amplitude (Petersen and Buzsáki, 2020). We hypothesize that this oscillatory adaptive mechanism serves to stabilize neural processing and prevent disruptive synaptic changes during speed variations.

### Phase coding across brain regions and species

Our findings, though focused on the hippocampus, extend to other brain regions where phase coding has been observed, such as the primary visual cortex and olfactory bulb. We have shown that in these areas, low-frequency oscillations—particularly in the high theta to low alpha range—also maximize information rate. Therefore, our results suggest a broader applicability of the speed-precision trade-off framework and encourage exploration of how single-cell properties constrain optimal frequency and noise levels across different brain areas and species. For instance, the two distinct theta rhythms in the human hippocampus (Goyal et al., 2020) might reflect different pyramidal neuron types with unique membrane properties tuned to process specific input characteristics. Hence, our findings offer a method to map single-cell properties to frequencies that optimize phase-coded information rate, providing a framework to assess the plausibility of such coding schemes across brain circuits.

### Applications to neuromorphic computing and artificial neural networks

Recent advancements in training spiking neural networks akin to standard artificial neural networks (Eshraghian et al., 2023; Neftci, Mostafa, and Zenke, 2019) have paved the way for optimizing large-scale spiking circuits for complex cognitive tasks, such as language production (Zhu, Zhao, et al., 2023) and autonomous driving (Zhu, Wang, et al., 2024). Incorporating efficient encoding strategies like phase coding, by using layer-wise reference oscillations or directly via complex-valued computations (Bybee, Frady, and Sommer, 2022), could significantly enhance the large-scale training of deep spiking networks. These sparse encoding formats would also support energy-efficient neuromorphic chips, enabling inference and training at the edge (Sun et al., 2024). Additionally, standard artificial neural networks—such as convolutional and recurrent neural networks—have been shown to benefit from reference oscillations in tasks related to visual perception (Duecker et al., 2023) and working memory (Pals, Macke, and Barak, 2024), suggesting promising directions for future research.

### Limitations and future work

Our theoretical framework relies on several assumptions that introduce limitations. One key assumption is the use of a noiseless model to estimate the expected spike phase, which does not account for the shift to earlier phases caused by high noise due to the asymmetry of spike threshold crossing. Additionally, by treating phase as a non-circular variable, we introduce a cutoff at *ϕ* = 0 (i.e., oscillation trough), resulting in boundary effects in simulations, where strong inputs under high noise cluster near *ϕ* = 0 (see Figure S2a, bottom-left panels, yellow distributions). Furthermore, neurons can produce multiple spikes per cycle in simulations, leading to cycle-to-cycle interference at certain frequencies (e.g., see Figure S2a, 10 Hz column, yellow distribution). Lastly, our estimation of phase variance is based on a linear approximation of system dynamics at threshold, which does not fully capture the underlying nonlinear dynamics. While this approximation is adequate for physiologically relevant noise levels (Figure 2b), it diverges from simulations at higher noise levels (see Figure S2b, and c). Although this divergence is largely due to phase being bounded to the range of 0 to 2*π* in simulations but unbounded in theory, the abovementioned factors limit the applicability of our framework, possibly requiring numerical validation when medium-to-high noise levels are considered. Furthermore, our framework does not explicitly address other significant factors, such as metabolic cost (e.g., bit/spike efficiency), the possibility of population coding strategies (e.g., potentially reducing variance as 1*/N* via sample mean decoding), and their interactions: increasing neuron count would enhance precision but also raise metabolic costs, suggesting optimal population sizes. Therefore, future work should consider these aspects to better understand how they influence optimal sampling frequencies in phase coding neurons under realistic metabolic conditions.

## Code Availability

The code and data to reproduce all the Figures can be found at https://github.com/adriamilcar/Phase-Coding-in-LIF.

## Acknowledgements

This work was supported by a HORIZON-EIC-2021 Pathfinder Challenges grant to CAVAA (project number 101071178).

## 5 Appendix

### 5.1 Full derivation of the mean phase of firing

To determine the expected spike phase, we focus on the deterministic part of the neuron model, assuming small noise amplitudes and Gaussian distributions. The general solution for an equation of the form:

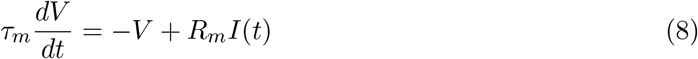

is given by:

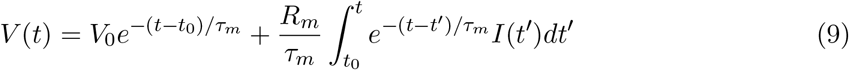

For the current *I*(*t*′) = *I*_*s*_ − *I*_*osc*_ cos (*ωt*′ + *ϕ*) in Equation (1), we compute two integrals. The first integral is:

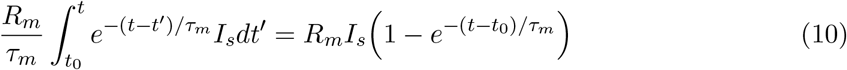

To handle the sinusoidal component, we use the fact that Re[∫ *f* (*x*)*dx*] = ∫ Re[*f* (*x*)]*dx*:

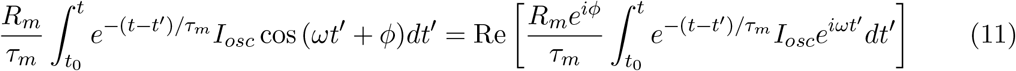

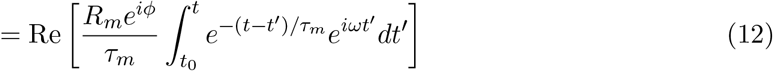

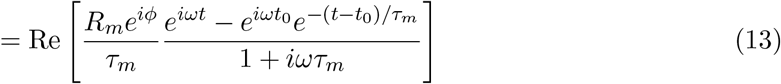

We factor out and express in polar coordinates:

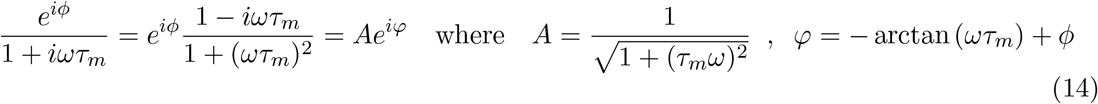

The real part of the expression is:

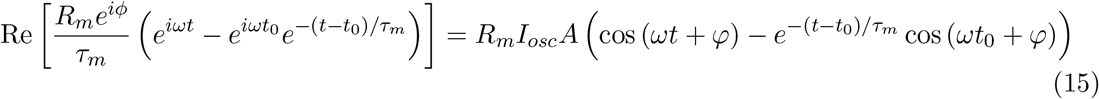

Substituting Equations (10) and (15) into Equation (9), we obtain the membrane potential:

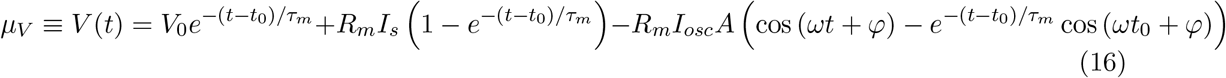

Then, assuming *V*_0_ = 0, the voltage trajectory, given a spike at time *t*′ and initial phase *ϕ*_0_, can be found as:

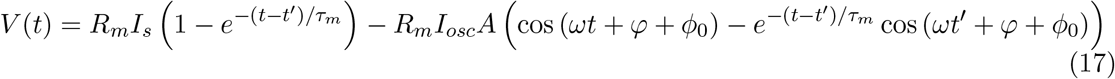

The spike phase of a neuron in cycle *i* is defined as the modulo 2*π* of the time of the first spike within the cycle 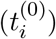, multiplied by the angular frequency *ω*:

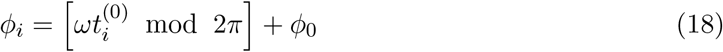

Imposing 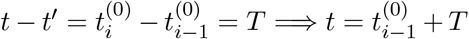, and substituting 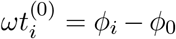 in Equation (17), we obtain:

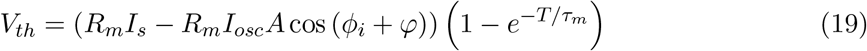

Isolating *ϕ*_*i*_, we get:

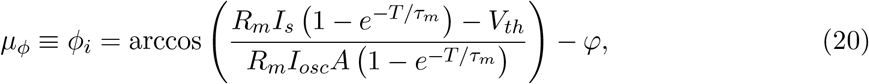

which gives the expected phase *µ*_*ϕ*_ at which the neurons will lock given an *I*_*s*_ value, replicating the main result in McLelland and Paulsen, 2009. Importantly, this equation constraints the range of *I*_*s*_ where the neuron will be within the phase-locking regime. To determine such a range, one must find the domain of the function, i.e., where −1 ≤ arccos ≤ 1 holds. Solving these inequalities, we obtain:

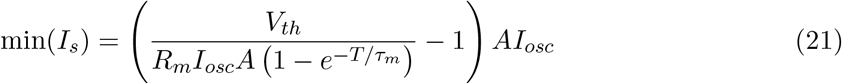

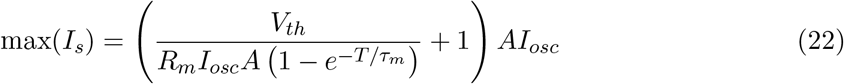

For simulations, we select *M* equally-spaced values between minimum and maximum, leaving a fraction (20%) of unused range on both extremes to alleviate boundary effects.

#### 5.2 Full derivation of the variance of phase of firing

To estimate the phase variance 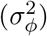 due to neuronal noise, we consider the variance in spike timing 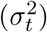 induced by fluctuations in the membrane potential (see Figure 3a). We employ a first-order Taylor series expansion and the propagation of uncertainty to derive an analytical approximation, along the lines of previous work (Demir and Sangiovanni-Vincentelli, 2012; Kilinc and Demir, 2018).

The membrane potential *V* (*t*) is approximated linearly around the spike threshold *V*_*th*_. We represent the membrane potential as *V* (*t*) = *V*_*th*_ + *δV*, where *δV* is a small deviation from the threshold. Using a Taylor series expansion, the potential at a slightly later time *t* + *δt* is given by:

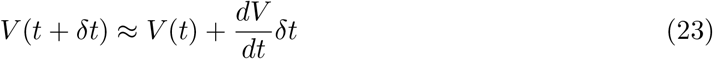

Since *V* (*t* + *δt*) − *V* (*t*) = *δV*, by rearranging the terms, we find the deviation in time as a function of the deviation in potential:

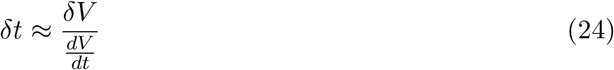

The variance in spike timing, 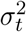, is related to the fluctuations in membrane potential (*δV*). Using the propagation of uncertainty, we express 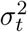 as:

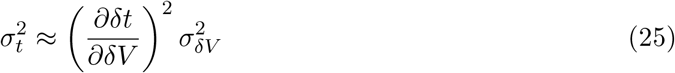

Substituting 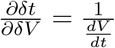 and recognizing that 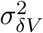 is the variance in membrane potential 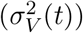, which is given by:

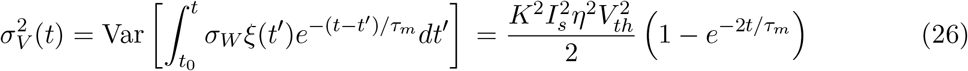

we obtain:

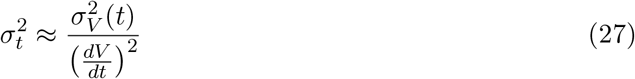

Then, we evaluate the expression at the threshold, so that:

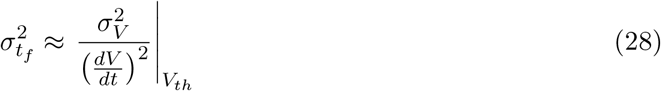

This approximation is valid under conditions where deviations from the spike threshold are small and the response of the membrane potential near the threshold is approximately linear. Finally, since time and phase are related by *ϕ* = *ωt*, we have that:

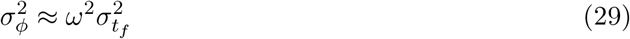

and under the assumption that the time of firing *t*_*f*_ will be the expected in the phase-locking regime 𝔼[*t*_*f*_] = *T*, we arrive at the expression:

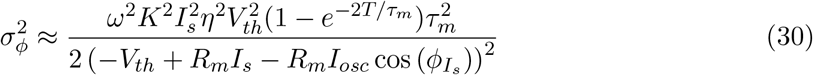

Equation 30 captures how the variance in the phase-of-firing is affected by the accumulation of random fluctuations in the membrane potential 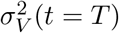, and the membrane potential dynamics around the threshold 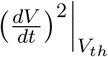. Then, since we are assuming small noise amplitudes in the suprathreshold regime (i.e., phase-locking regime) and given that we use linear approximations, the phase probability distribution can be approximated by a Gaussian:

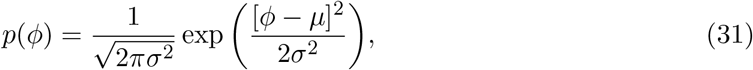

with mean 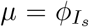, given by Equation 3, and variance 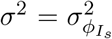, given by Equation 30, for a certain tonic input *I*_*s*_.

#### 5.3 Approximation of information rate

We first define the phase as the response *R* associated with a stimulus *S*, corresponding to a tonic input current *I*_*s*_. Assuming *M* tonic input levels that are equally likely, *S* is defined as a uniformly distributed variable *p*(*S* = *s*) = 1*/M*. In turn, the unconditioned distribution of *R* corresponds to a Gaussian mixture with equal weights:

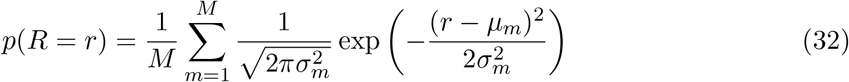

Here, *µ*_*m*_ and 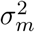 are the means and variances associated with each tonic input level *I*_*s*_ (i.e., the Gaussian components). In a Gaussian mixture model, the mutual information I(*S*; *R*) between *S* and *R* is given by the difference between the entropy of the Gaussian mixture (unconditioned response) and the average entropy of the individual Gaussian components (response conditioned to the stimulus):

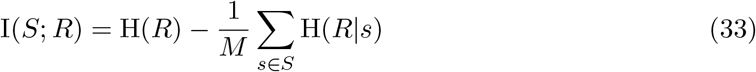

The entropy of a Gaussian mixture H(*R*) does not have a known closed-form solution (Huber et al., 2008). However, we have found that H(*R*) can be well approximated by considering an aggregate measure of the overall spread of the mixture: combining the variance of means ***µ*** and the mean of variances ***σ***^2^, such that:

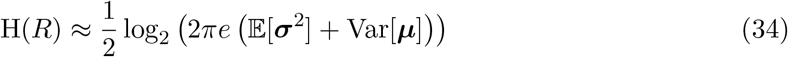

Then, the average entropy of the individual Gaussian components, H(*R*|*S*), does have a closedform solution:

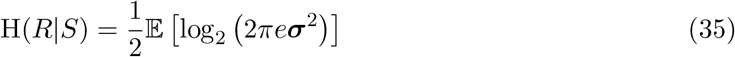

Combining these, and simplifying, we obtain:

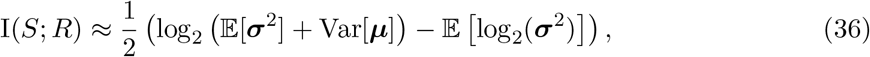

providing a good approximation of the average information of the phase of the first spike in one cycle about the input signal conveyed by *I*_*s*_.

One could then estimate the information rate *r* simply by multiplying the information per cycle I and sampling frequency *f*, such that r ≈ I*f*. However, this approach assumes statistical independence between cycles, providing only an upper bound on r. Instead, a more realistic approach would be to consider that the oscillation is sampling from an input signal *s* that has a characteristic time constant *τ*_*s*_, following an exponential decay in its autocorrelation function 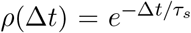. Then, the total correlation between one cycle and all subsequent cycles can be represented as a geometric series:

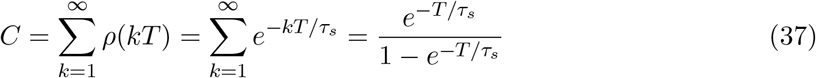

The sum *C* represents the cumulative effect of correlations across all cycles. Thus, we define the effective sampling frequency as 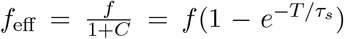, with 1 + *C* correcting the information rate by accounting for redundancy, and keeping *f*_eff_ = 0 as *C* → ∞ and *f*_eff_ = *f* as *C* → 0. After simplifying, we get the expression:

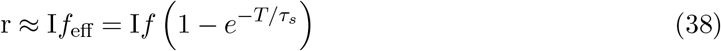

This expression ensures that the information rate reflects the impact of temporal correlations in the sampled signal, penalizing for oversampling when *T* < *τ*_*s*_.

Finally, to find the optimal frequency for a particular set of neuron parameters and noise level, we define the normalized information rate (r_*norm*_) as the ratio of r at a specific frequency (r_*f*_) to the sum of r across all frequencies (*F*):

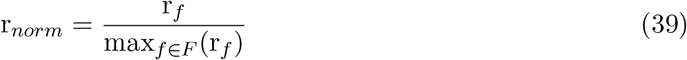

The optimal frequency *f*\* is then defined as the frequency at which r_*norm*_ is maximized, indicating the frequency that provides the highest information rate:

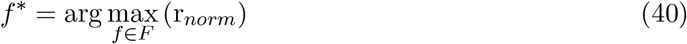

This optimal frequency *f*\* will be determined by the trade-off between sampling speed and encoding accuracy imposed by noise.

#### 5.4 Neuron parameters

By default, and unless it is stated otherwise in the main text, we used the parameters shown in Table 1. For the neuron and oscillation parameters, we followed McLelland and Paulsen (2009) and for the noise strength Lansky, Sanda, and He (2006), setting 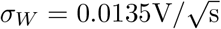.

**Table 1:** Default parameters for hippocampal neurons.

| Parameter | Description | Value |
| --- | --- | --- |
| $R_m$ | Membrane resistance | 142 MΩ |
| $V_{th}$ | Spike threshold | 15 mV |
| $\tau_m$ | Membrane time constant | 24 ms |
| $I_{osc}$ | Oscillation amplitude | 40 pA |
| $\sigma_W$ | Noise amplitude | $0.0135 \text{ V}/\sqrt{\text{s}}$ |

For the parameter gradients along the dorsoventral axis, we followed Malik et al. (2016) and Patel et al. (2012). For the noise strength, we followed Lansky, Sanda, and He (2006), setting 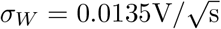 across models. The neuron parameters can be found in Table 2.

**Table 2:**
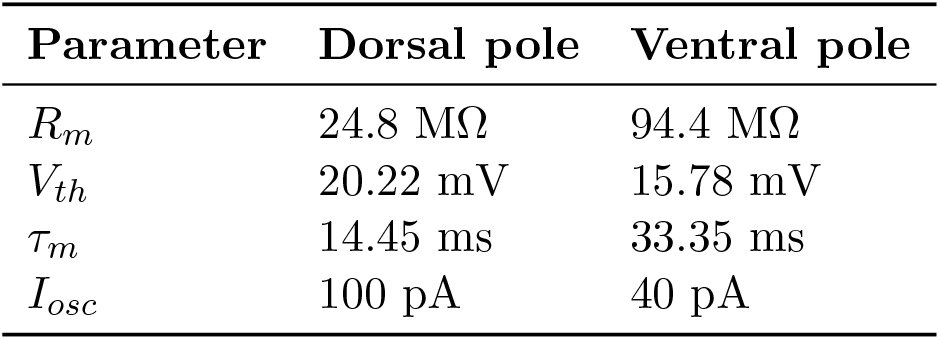
Neuron parameters along the hippocampal dorsoventral axis.

| Parameter | Dorsal pole | Ventral pole |
| --- | --- | --- |
| $R_m$ | 24.8 MΩ | 94.4 MΩ |
| $V_{th}$ | 20.22 mV | 15.78 mV |
| $\tau_m$ | 14.45 ms | 33.35 ms |
| $I_{osc}$ | 100 pA | 40 pA |

We based the parameters for pyramidal cells in the primary visual cortex on Teeter et al. (2018). For mitral cells in the olfactory bulb, we followed Margrie and Schaefer (2003), using an oscillatory current of *I*_*osc*_ = 35.4 pA to produce realistic membrane potential amplitudes of 3.45 mV in response to a 3.7 Hz input oscillation. The list of parameters is provided in Table 3.

**Table 3:**
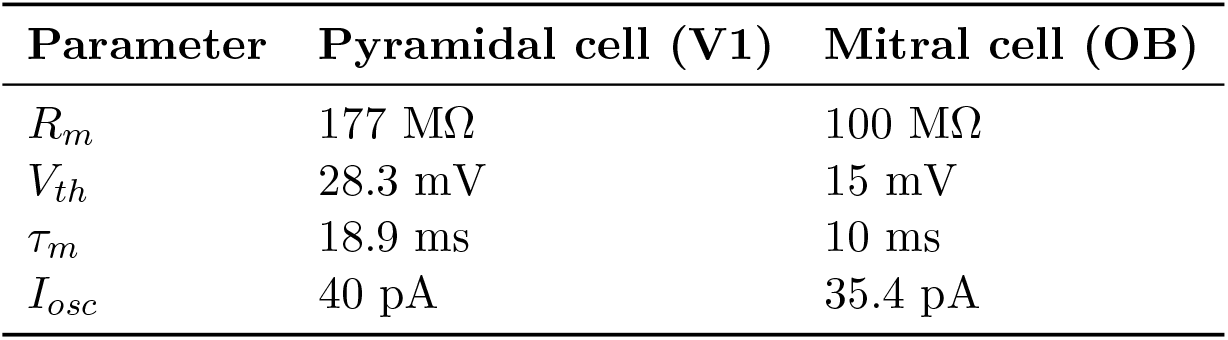
Neuron parameters for visual and olfactory cells.

| Parameter | Pyramidal cell (V1) | Mitral cell (OB) |
| --- | --- | --- |
| $R_m$ | 177 MΩ | 100 MΩ |
| $V_{th}$ | 28.3 mV | 15 mV |
| $\tau_m$ | 18.9 ms | 10 ms |
| $I_{osc}$ | 40 pA | 35.4 pA |

### 6 Supplementary figures

**Figure S1:**
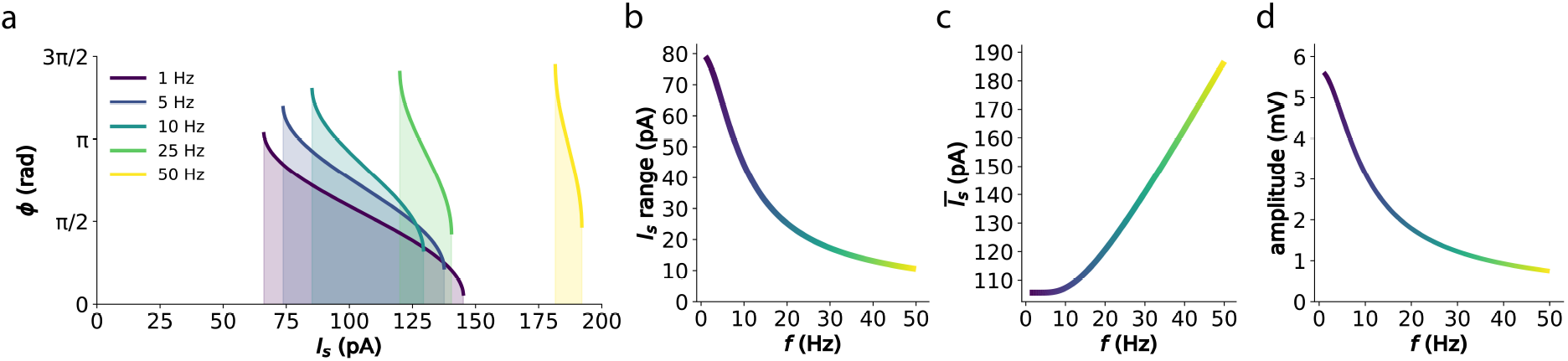
Oscillatory frequency modulates the tonic input range that makes neurons phase-lock. (a) Phase-locking function for different values of the oscillatory frequency *f* (color coded). Shaded areas denote the phase-locking range of *I*_*s*_, corresponding to the domain of *I*_*s*_ in Equation 3. The parameters are the same ones used in McLelland and Paulsen (2009) and in Figure 1, to match hippocampal physiology. (b) Length of *I*_*s*_ range (max(*I*_*s*_) − min(*I*_*s*_) = 2*AI*_*osc*_) across frequencies. (c) Average *I*_*s*_ as 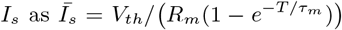 (middle point in *I*_*s*_ range) across frequencies. (d) Effective amplitude of the membrane potential oscillation, *V*_*osc*_ produced by the oscillatory input *I*_*osc*_. Given that the membrane acts as a low-pass filter, determined by *τ*_*m*_, the effective oscillation in the membrane potential can be found to be *V*_*osc*_ = *R*_*m*_*I*_*osc*_*A*. Thus, since the membrane filters the oscillatory input as 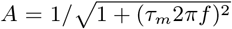, the amplitude of the membrane potential *V*_*osc*_ will decrease with *f* approximately as *~* 1*/f*.

**Figure S2:**
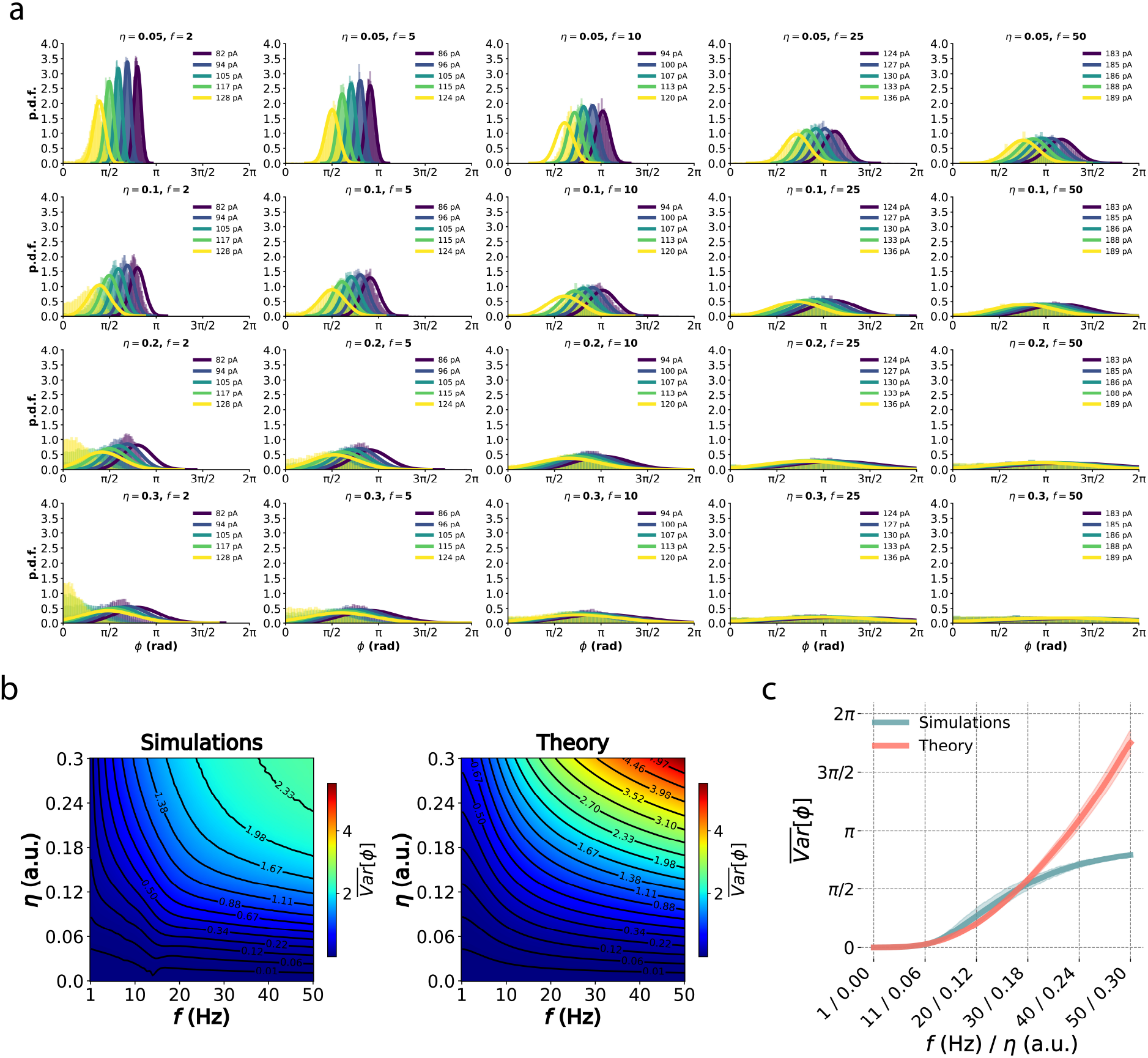
Analytical approximation of phase distributions. (a) Phase distributions for a range of frequencies and noise strengths. Histograms denote the simulations whereas solid lines denote the theoretical predictions. For the simulations, first-spike phases are recorded from the beginning of the second cycle (with the trough as *ϕ* = 0), after initializing the neurons to their expected phase *ϕ*_0_ = *µ*_*ϕ*_ to allow them to reach steady-state dynamics. (b) Average variance in rad^2^ (across *I*_*s*_ levels) across a wide frequency-noise parameter space, for simulations and the theoretical predictions. (c) Diagonal slices of plots in (b), showing the deviation of the theory from the simulations after a certain level of noise amplitudes at high frequencies, due to the bounded variance of simulated spike phases constrained to the measurable range of [0, 2*π*] radians.

**Figure S3:**
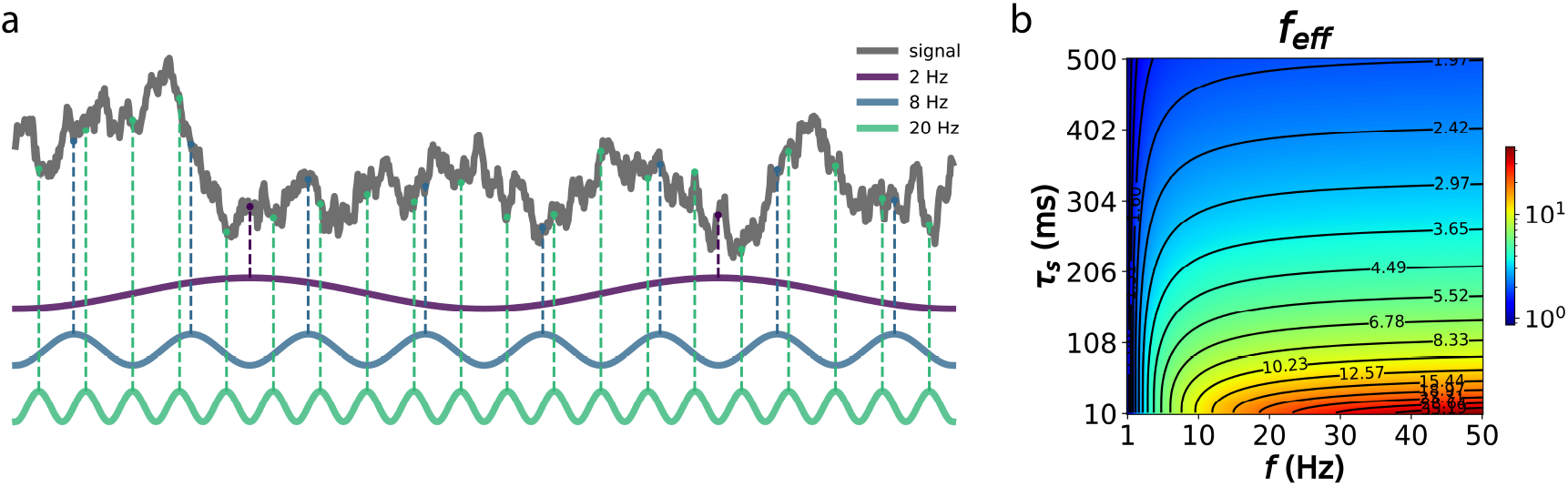
Effective rhythmic input sampling. (a) An example signal with *τ*_*s*_ of 100 ms sampled by different oscillation frequencies. (b) Effective frequency 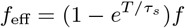 for various *τ*_*s*_ values.

**Figure S4:**
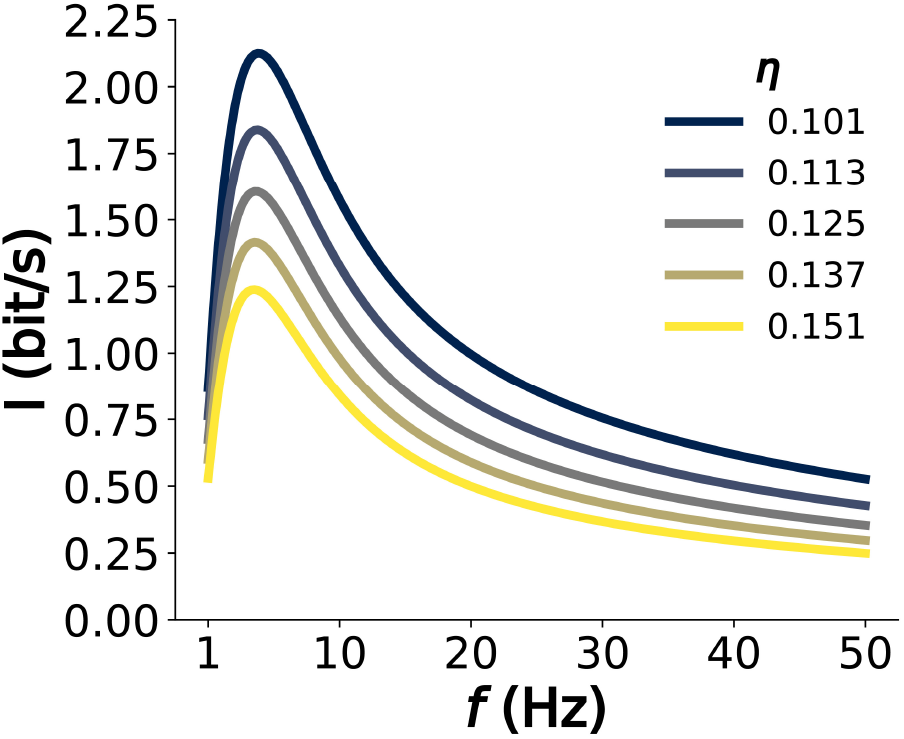
Information rate across frequencies for the range of physiologically realistic noise levels (*η*).

**Figure S5:**
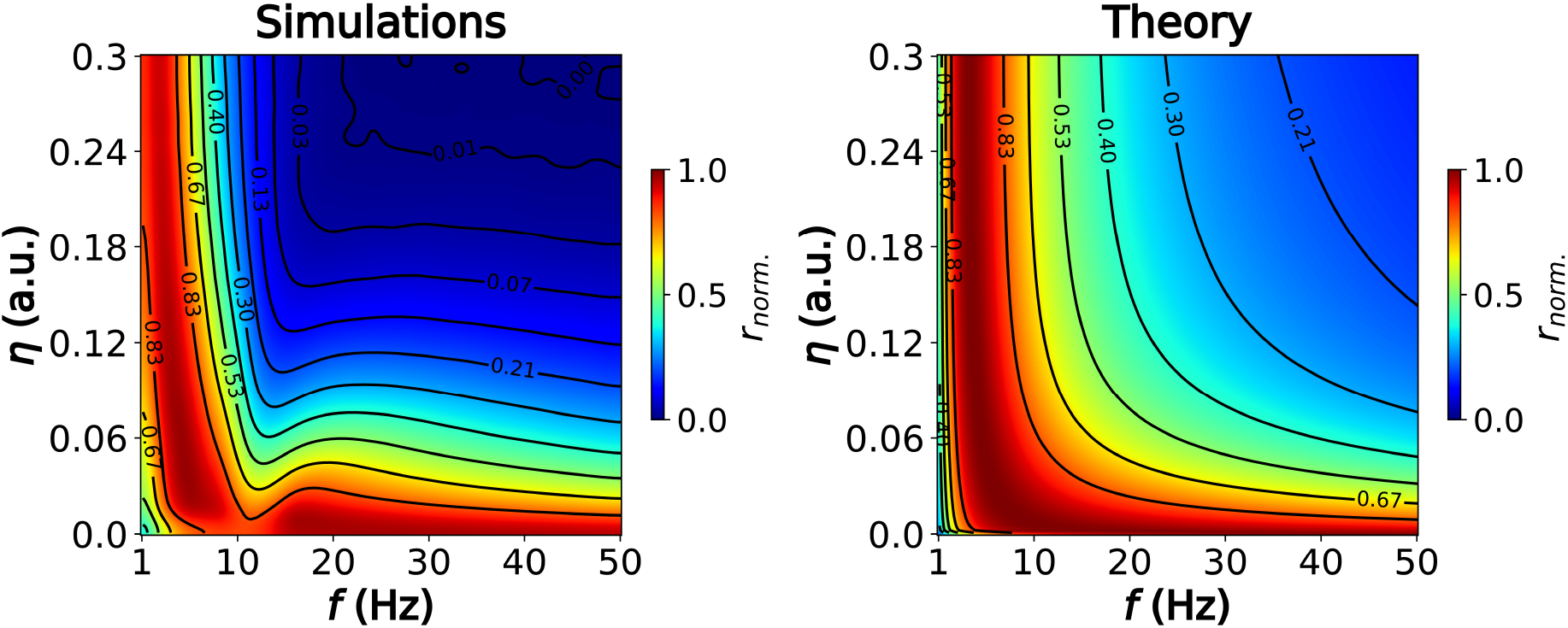
Normalized information rate across the frequency-noise parameter space for simulations and theoretical predictions.

**Figure S6:**
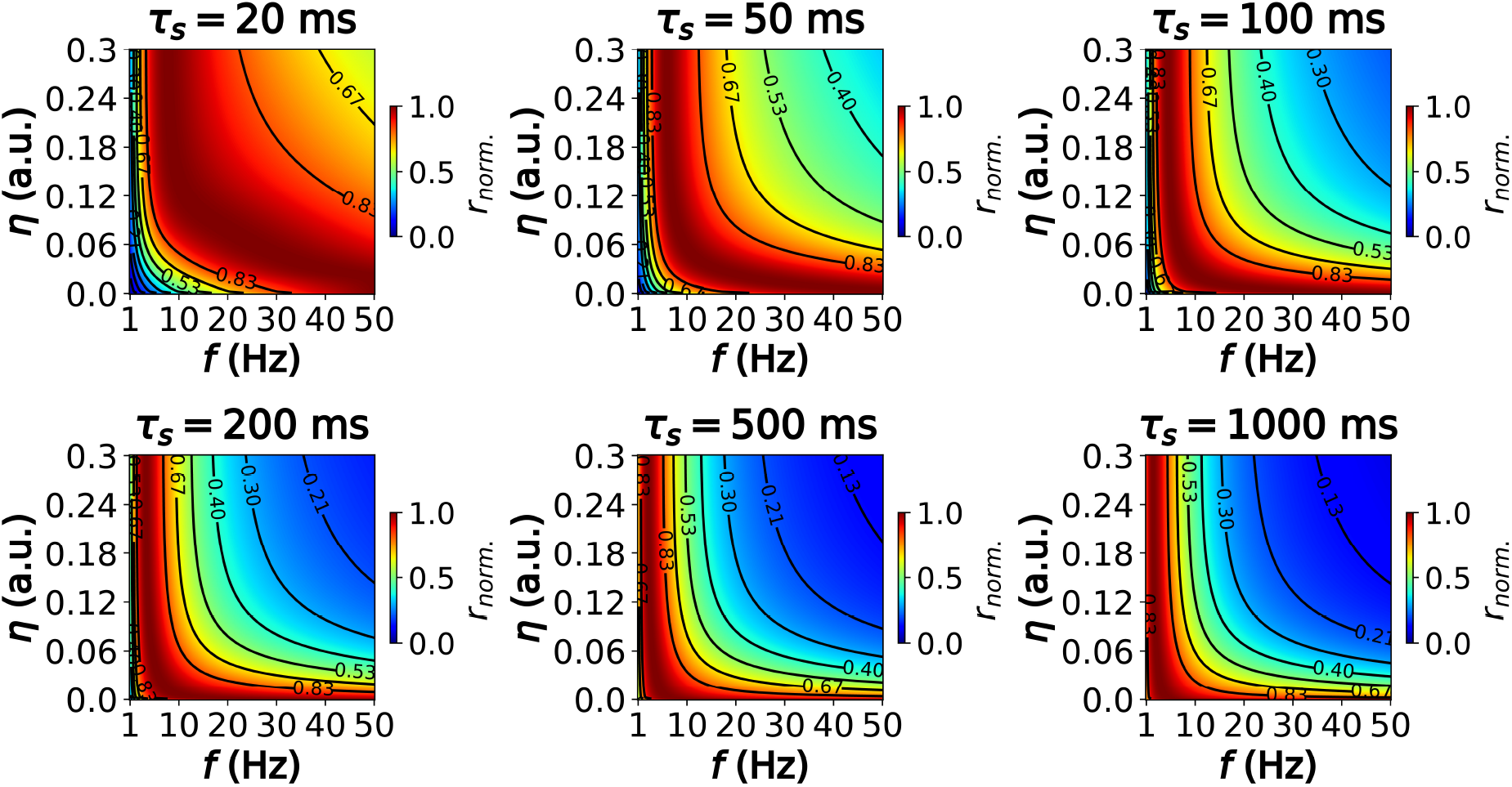
Normalized information rate across the frequency-noise parameter space for a wide range of input signal time constants *τ*_*s*_.

**Figure S7:**
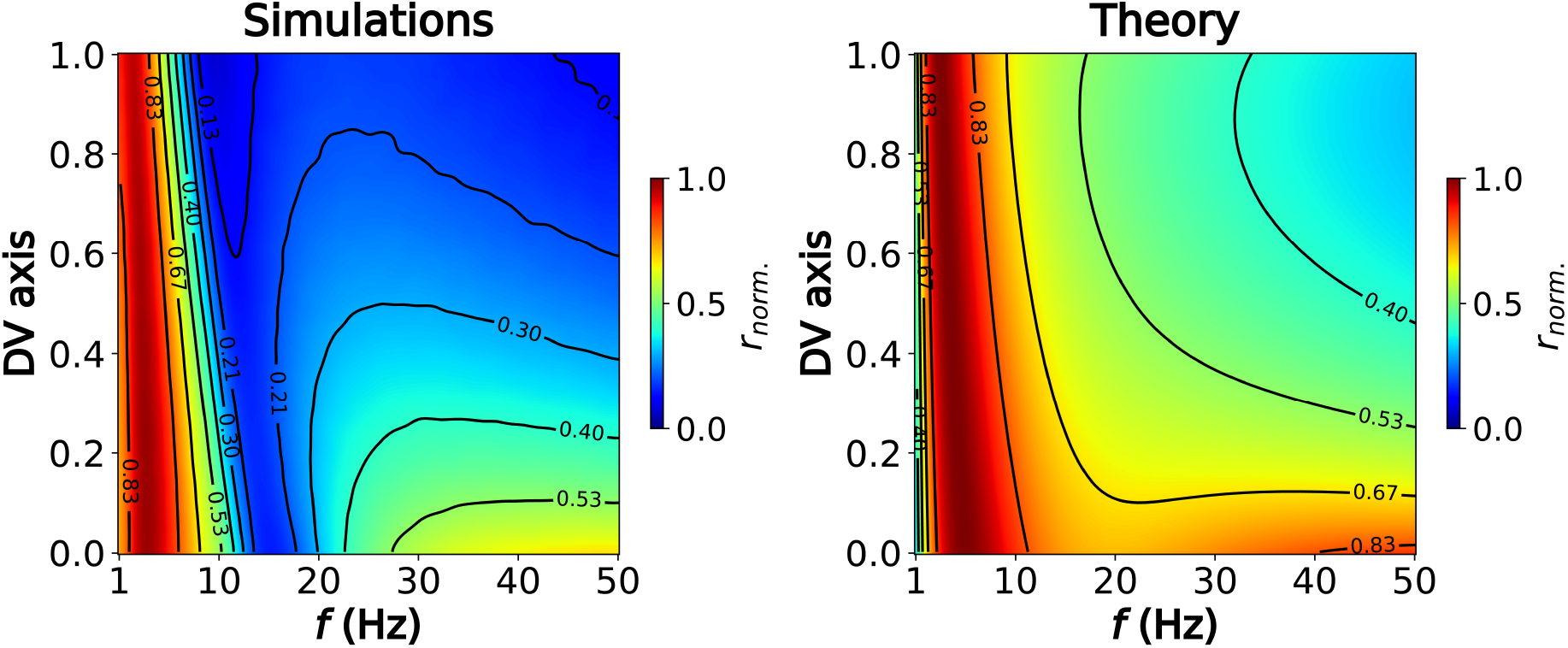
Normalized information rate across the dorsoventral axis for simulations and theoretical predictions.

**Figure S8:**
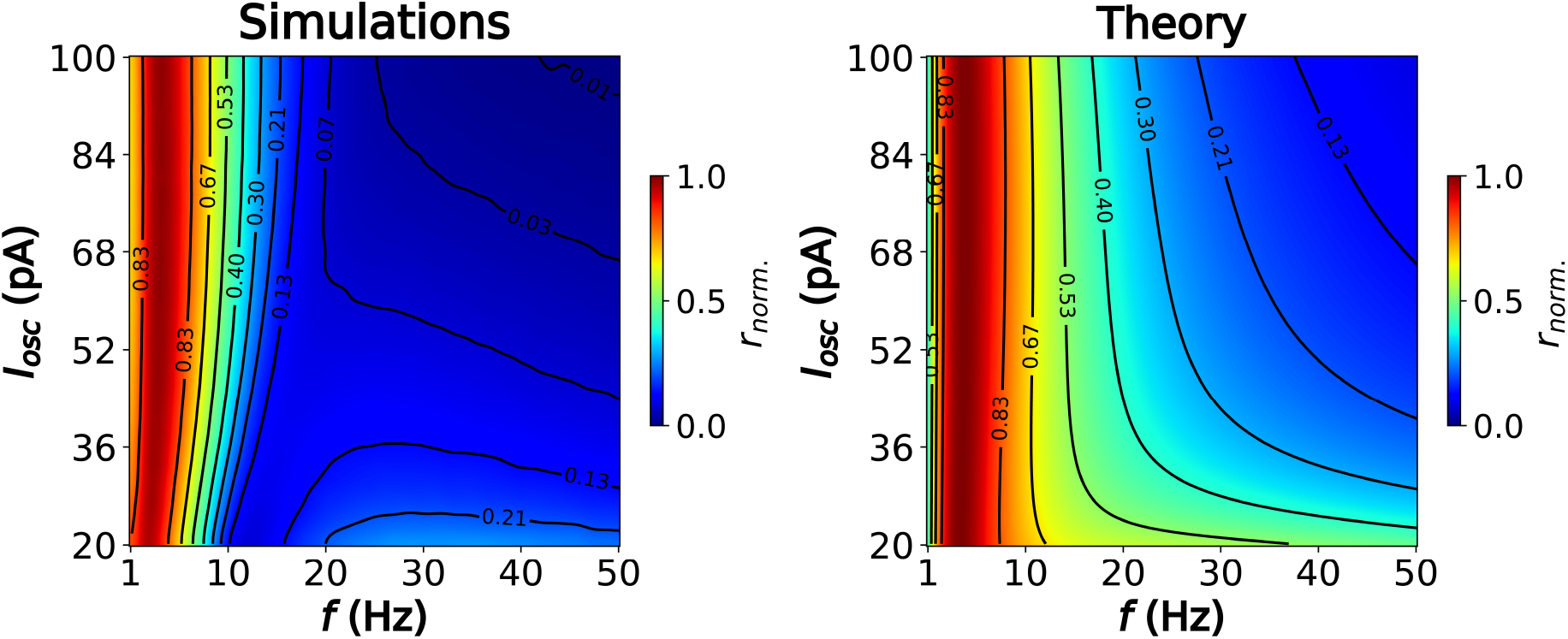
Normalized information rate across the frequency-amplitude space for simulations and theoretical predictions.

## Notes

https://github.com/adriamilcar/Phase-Coding-in-LIF

### Competing Interest Statement

The authors have declared no competing interest.

### Summary of Updates

Equations 5,6,7 have been updated to provide with a proper approximation of the information rate in phase coding neurons (and not just a proportional approximation, as it was before). Then, Figure 3 has been updated accordingly. In addition, a supplementary figure (now S4) has been added to report the actual information rate (bits/s) that is estimated for physiologically realistic levels of noise in the brain. Some minor mistakes have been corrected in Appendix 5.1. The paragraph on Noise (pages 3-4) and the corresponding Equation 2 have been updated to improve clarity. Finally, we added a few more citations in the Discussion to better acknowledge previous and recent work in the field.

